# Neutrophil extracellular traps promote tumor chemoresistance to anthracyclines

**DOI:** 10.1101/2024.11.07.622533

**Authors:** Cindy Lin, Sarah E. Herlihy, Marina Li, Hui Deng, Rina Kim, Luca Bernabei, Matthew Rosenwasser, Dmitry I. Gabrilovich, Dan T. Vogl, Yulia Nefedova

## Abstract

The microenvironment plays an important role in promoting tumor cell chemoresistance, but the mechanisms responsible for this effect are not clear. Here, using models of multiple myeloma (MM) and solid cancers, we demonstrate a novel mechanism mediated by neutrophils, a major cell population in the bone marrow (BM), that protects cancer cells from chemotherapeutics. We show that in response to tumor-derived soluble factors, BM neutrophils release their DNA in the form of neutrophil extracellular traps (NETs). Cell-free DNA derived from NETs is then taken up by tumor cells via endocytosis and localizes to the cytoplasm. We found that both NETs and cell-free DNA taken up by tumor cells can bind anthracyclines, leading to tumor cell resistance to this class of chemotherapeutic agents. Targeting cell-free DNA with Pulmozyme or blocking NET formation with a PAD4 inhibitor abrogates the chemoprotective effect of neutrophils and restores sensitivity of tumor cells to anthracyclines.

## Introduction

Chemoresistance is a major clinical challenge in cancer treatment. The tumor microenvironment plays an important role in mediating resistance and survival of tumor cells and the eventual relapse and progression of malignant disease.

Neutophils comprise the largest cell population within the bone marrow (BM), but their role in progression of cancers that originate or metastasize to the BM remains largely unknown. Several recent reports indicated that an elevated neutrophil/lymphocyte ratio is associated with a poor prognosis in multiple myeloma (MM), a malignancy of plasma cells localized in the BM (Kelkitli et al., 2014; Li et al., 2017; Romano et al., 2015; Shi et al., 2017; Wongrakpanich et al., 2016). Our group recently demonstrated that neutrophils protect MM cells from chemotherapeutic agents, including anthracyclines (Ramachandran et al., 2016). However, the mechanism by which neutrophils mediate their chemoprotective effect remains unknown.

Neutrophil extracellular traps (NETs) were originally identified as a mechanism of binding and killing pathogens that are too large to be engulfed (Brinkmann et al., 2004). NETs are structures consisting of chromatin fibers (DNA and histones) and proteins, the majority of which are normally found in neutrophil cytoplasm or granules (Urban et al., 2009). NETs are generated by dying neutrophils in a process termed NETosis (Fuchs et al., 2007; Wang et al., 2009). However, under certain conditions NETs can also be formed solely from mitochondrial (mt) DNA, in which case neutrophils remain viable after NET formation (Yousefi et al., 2009). Formation of NETs is a multi-step process that typically requires activity of peptidylarginine deiminase 4 (PAD4). PAD4 citrullinates histones, including histone H3, and thus promotes decondensation of chromatin, a necessary step during NETosis (Wang et al., 2009). NET formation has been shown to be triggered by a variety of proinflammatory stimuli, including reactive oxygen species, bacterial lipopolysaccharide, tumor necrosis factor alpha, and interleukin-8 (Brinkmann et al., 2004; Fuchs et al., 2007; Tadie et al., 2013).

Recently, NETs have been implicated in solid tumor progression. Neutrophils form NETs in cancer patients, in tumor-bearing (TB) animals, and *ex vivo* (reviewed in (Masucci et al., 2020). NETs promote mouse lung carcinoma and breast cancer cell trapping and metastasis (Cools-Lartigue et al., 2013; Park et al., 2016) and have been linked with tumor-associated thrombosis (Abdol Razak et al., 2017). Here, we report our unexpected finding that NETs, via their associated DNA, prevent the antitumor activity of the anthracycline class of chemotherapeutics and thus may contribute to chemotherapy resistance. It suggests a therapeutic approach to its regulation.

## Results

### Multiple myeloma cells induce NETs and cell-free DNA release by neutrophils

Very little is known about whether neutrophils from tumor-free (TF) and TB hosts have a similar ability to form NETs in response to tumor-derived factors. To address this question, we isolated CD11b^+^Ly6G^+^ cells from BM of TF or DP42 MM-bearing mice. We and others previously demonstrated that the majority of neutrophils in TB mice are pathologically activated immune suppressive polymorphonuclear myeloid-derived suppressor cells (PMN-MDSC)(Ramachandran et al., 2013). In this study, we did not specifically evaluate the functionality of these cells, therefore we referred to all cells as neutrophils. Cells were cultured *in vitro* with DP42 MM cells separated by Transwell inserts. Unstimulated CD11b^+^Ly6G^+^ cells from TF and TB mice had low ability to spontaneously form NETs. The presence of MM cells led to a significant increase in NET formation by neutrophils from both TB and TF mice; however, the amount of NET formation by neutrophils from TB mice was substantially higher than by neutrophils from TF mice (**Figure 1A**). Similar data was obtained using BM neutrophils isolated from mice bearing orthotopic AT3 breast cancer (**Figure 1B**). Formation of NETs was accompanied by a significant increase in the level of cell-free DNA (cfDNA, **Figure 1C**) and, specifically, cfDNA bound to citrullinated histone H3 (**Figure 1D**). Changes in citrullinated histone H3-bound cfDNA reflected changes in NETosis (**Figure 1A and 1D**).

**Figure 1.**
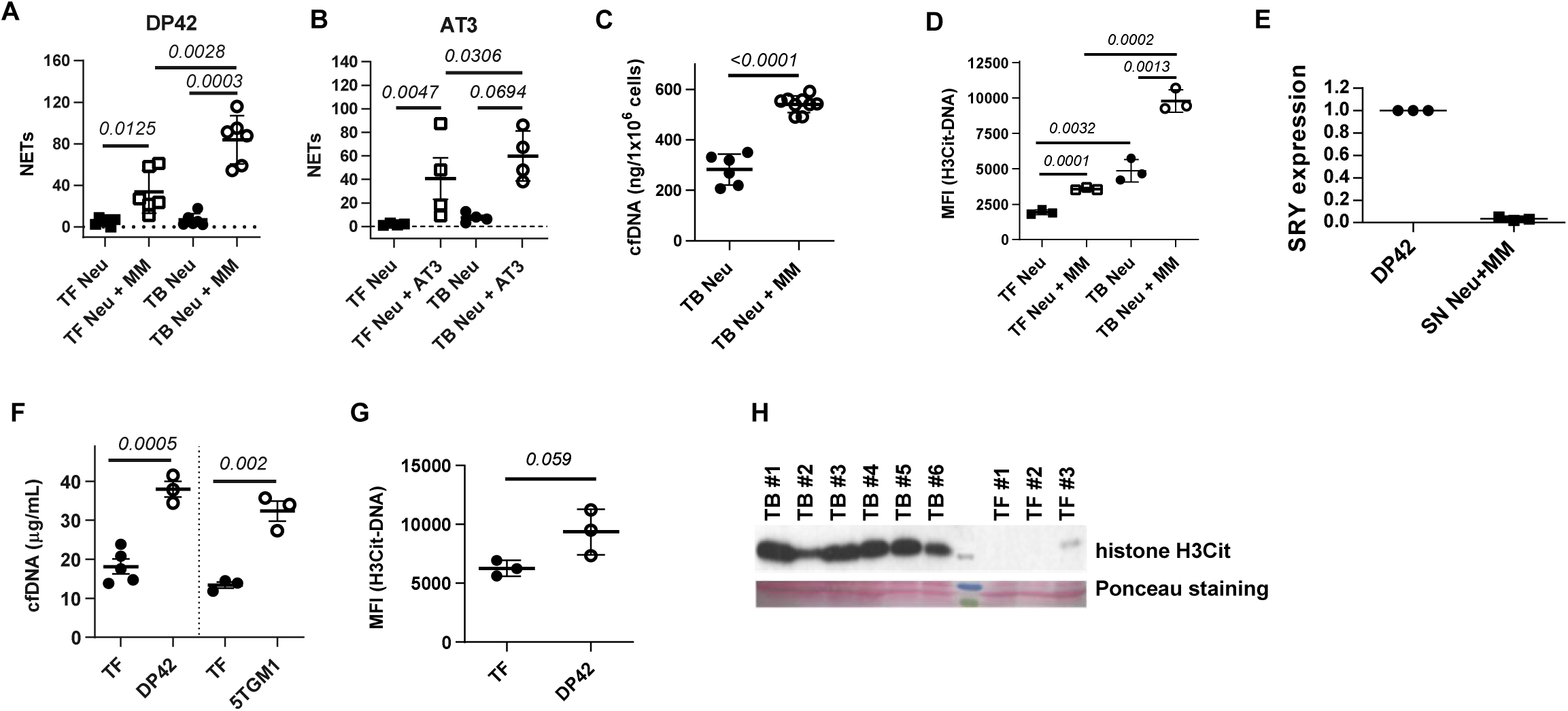
Mouse MM induces NETosis. (**A,B**) Formation of NETs by mouse BM neutrophils (Neu) isolated from control TF or TB mice and cultured in the presence or absence of mouse MM DP42 cells (**A**) or mouse AT3 breast cancer cells (**B**) separated by Transwell insert. Quantitation of NET area per cell (μm^2^/cell) is shown. (**C,D**) Concentration of cfDNA (**C**) or cfDNA bound to citrullinated histone H3 (**D**) in supernatants from neutrophils isolated from mouse BM and cultured alone or in the presence of DP42 MM cells separated by Transwell insert. (**E**) Gene expression of SRY was determined by real-time PCR using genomic DNA isolated from DP42 cells (DP42) and cfDNA isolated from neutrophil-MM cell culture supernatants (SN Neu+MM). (**F,G**) Level of cfDNA (**F**) or cfDNA bound to citrullinated histone H3 (**G**) measured in the BM of TF mice and indicated TB mice. (**H**) Presence of citrullinated histone H3 was detected in BM flush collected from TF and DP42 TB mice. Statistics: two-tailed Student’s *t* test.

We hypothesized that NETs represent a major source of cfDNA. To determine the cellular origin of cfDNA, neutrophils from female mice were co-cultured with MM cells originally derived from male mice. PCR of the sex-determining region of the Y chromosome of the cfDNA isolated from supernatants of neutrophil-MM cell co-cultures demonstrated that neutrophils, but not MM cells, were the main source of cfDNA (**Figure 1E**). Release of DNA was associated with death of neutrophils from both TF and TB mice (**Figure S1A**), but not with increased apoptosis (**Figure S1B**), which supports the conclusion that the cfDNA originates from NETs. Growth of MM in the syngeneic mouse models DP42 and 5TGM1 was accompanied by a marked increase in the amount of total cfDNA in the BM (**Figure 1F**), cfDNA bound to citrullinated histone H3 (**Figure 1G**) and citrullinated histone H3 (**Figure 1H**) compared to control TF mice.

We investigated which soluble factor(s) may be responsible for MM-induced NETosis. CD138 (Syndecan-1, SDC1) is highly expressed by MM cells and is known to be shed by these cells. High mobility group box protein 1 (HMGB1), normally an intracellular protein, is known to be released by solid tumor cells. We demonstrated the presence of HMGB1 protein in supernatants collected from a panel of MM cells (**Figure S2**). Both SDC1 and HMGB1 induced NET formation at levels comparable to that induced by MM cells (**Figure 2A**). Addition of anti-HMGB1 neutralizing antibodies (**Figure 2B**) or pharmacological inhibition of SDC1 shedding (**Figure 2C**) abrogated MM-induced NET formation. TLR4 and RAGE have been previously reported to serve as the main receptors for HMGB1 (Li et al., 2013). We found no differences in NETs induced by MM cells between BM neutrophils isolated from RAGE KO and WT mice. However, MM cells as well as recombinant HMGB1 or SDC1 were unable to induce NETosis in neutrophils isolated from TLR4 KO mice (**Figure 2D**).

**Figure 2.**
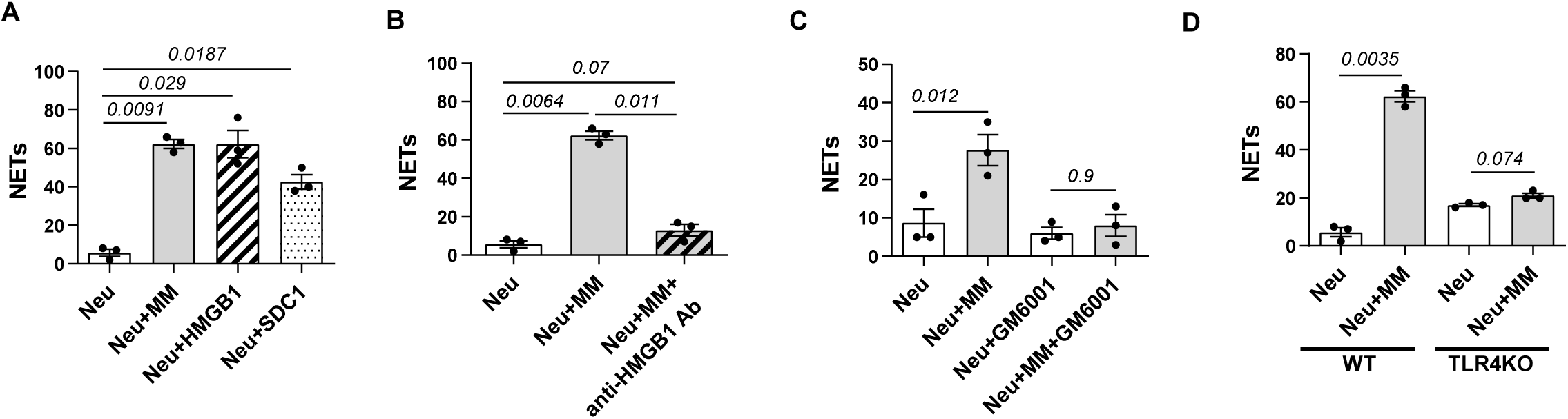
MM-induced NETosis is mediated through TLR4. Formation of NETs by mouse BM neutrophils cultured in the presence or absence of DP42 cells, 200 ng/mL HMGB1, or 5 µg/mL SDC1 (A), with or without anti-HMGB1 neutralizing Ab (5 µg/mL) (B), GM6001 (10 µM) (C) or by neutrophils isolated from TLR4 KO mice (D) was evaluated by microscopy and quantitated. Individual values, mean, and SD are shown. Statistics: One-way ANOVA (A-C), two-tailed Student’s *t* test (D).

Next, we tested the effect of human tumor cells on NET formation by BM neutrophils. MM cells induced NET formation by neutrophils isolated from both healthy donors and MM patients (**Figure 3A-D and Figure S3A**). As anticipated, analysis of MM-induced NETs revealed the presence of citrullinated histone H3 (**Figure 3A**). BM neutrophils from MM patients had a greater ability to form NETs in response to human MM cell lines (HMCLs) than neutrophils from donors (5.7±2.3 fold and 3.6±1.3 fold increase, respectively; **Figure 3B, C**). HMCLs and primary CD138^+^ MM cells isolated from BM of patients similarly induced NET formation (**Figure 3C**). Neutrophils isolated from peripheral blood (PB) of MM patients also underwent NETosis in response to HMCLs (**Figure 3D**); however, the ability of PB neutrophils to form NETs was significantly lower than that of BM neutrophils (1.4±0.1 fold and 5.7±2.3 fold, respectively). Formation of NETs by BM neutrophils from MM patients, either induced by HMCLs or primary MM cells, was accompanied by a significant increase in cfDNA (**Figure 3E**). Previous data indicated that NETs may contain either genomic DNA (gDNA) or mitochondrial DNA (mtDNA) (Yousefi et al., 2009). PCR analysis of cfDNA purified from neutrophil-MM supernatants demonstrated the presence of genes encoded by both types of DNA: gDNA (*FAS, GAPDH, RHOH, ACTB*) and mtDNA (*ND1, CO1, ATP6, CYB*) (**Figure S3B**).

**Figure 3.**
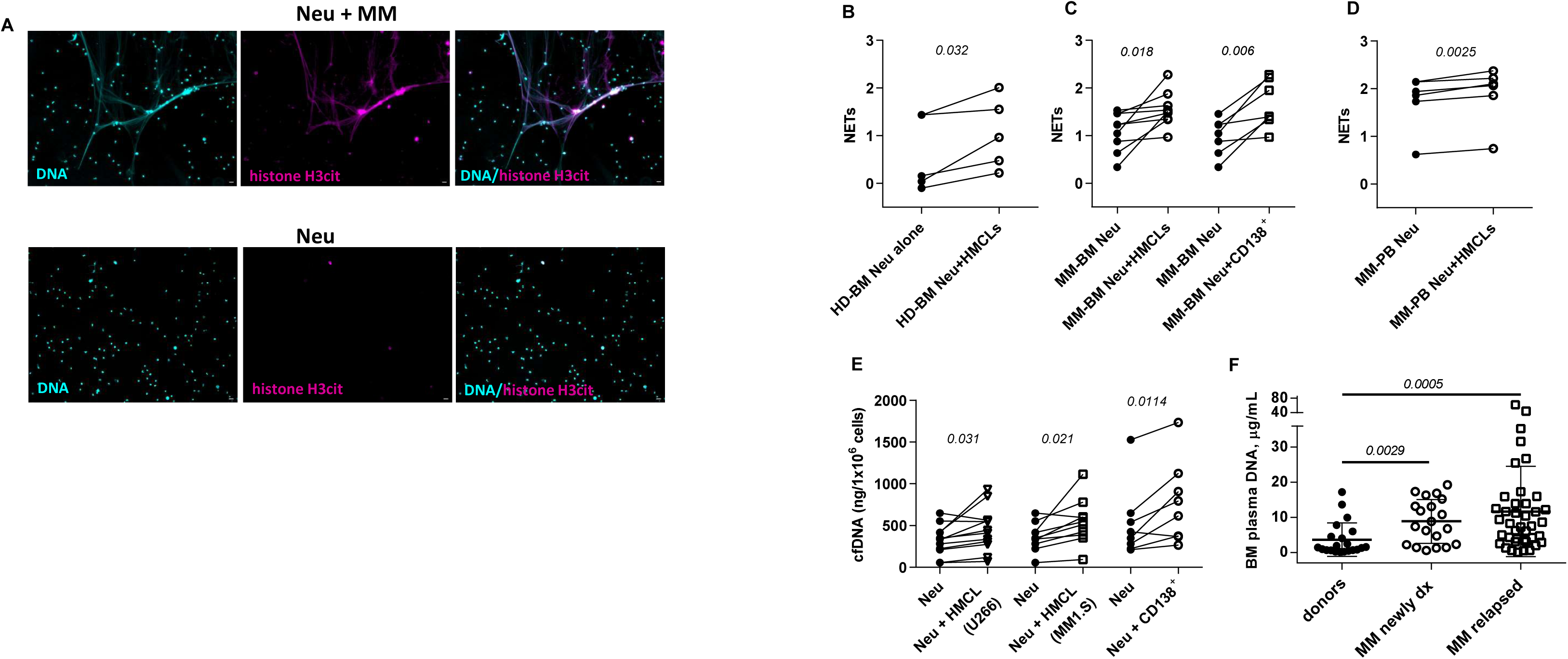
Human MM cells induce NETosis. (**A**) Formation of NETs in the presence or absence of human MM cells. Representative images of NET’s DNA co-localized with citrullinated histone H3. Magnification 20x; scale bar, 200 µm. (**B-D**) Formation of NETs by neutrophils isolated from the BM of (**B**) healthy donors (n=5), (**C**) MM patients (n=9), and (**D**) neutrophils from PB of MM patients (n=6) in the presence or absence of HMCLs or primary MM cells (CD138^+^). Data presented as log_10_ NET area (μm^2^/cell). Individual values are shown. Statistics: paired two-tailed Student’s *t* test. (**E**) Concentration of cfDNA measured in supernatants collected after overnight culture of neutrophils from MM patients’ BM, with or without the indicated MM cell lines (n=12) or CD138^+^ primary MM cells (n=7) separated by Transwell insert. Individual values are shown. Statistics: paired two-tailed Student’s *t* test. (**F**) Levels of DNA in BM plasma of healthy donors (n=21), newly diagnosed MM patients (n=20) and relapsed/refractory MM patients (n=40). Individual values, mean, and SD are shown. Statistics: Mann Whitney test.

To confirm the clinical relevance of these data, DNA concentration was measured in BM plasma obtained from newly diagnosed untreated MM patients, relapsed MM patients, and healthy donors. BM plasma cfDNA was significantly increased in MM patients compared to healthy donors. Relapsed MM patients had higher levels of BM cfDNA than newly diagnosed MM patients, however, the difference did not reach statistical significance (**Figure 3F**). There were no significant correlations between cfDNA levels and the proportion of plasma cells in the BM aspirate and core biopsy, levels of paraprotein, beta-2 microglobulin, karyotype and FISH abnormalities, or cell differential count in the BM or PB.

### cfDNA released by neutrophils neutralize the antitumor effect of doxorubicin

Our previous data demonstrated that neutrophils can protect MM cells from chemotherapy through a cell contact-independent mechanism (Ramachandran et al., 2016). We hypothesized that cfDNA is one of the factors responsible for this effect. To test this hypothesis, human MM cells were treated with doxorubicin in the presence of cfDNA purified from supernatants from human or mouse neutrophil-MM cultures. In parallel, gDNA extracted from mouse or human neutrophils was used. All tested DNA similarly protected MM cells from doxorubicin-induced apoptosis (**Figure 4A**).

**Figure 4.**
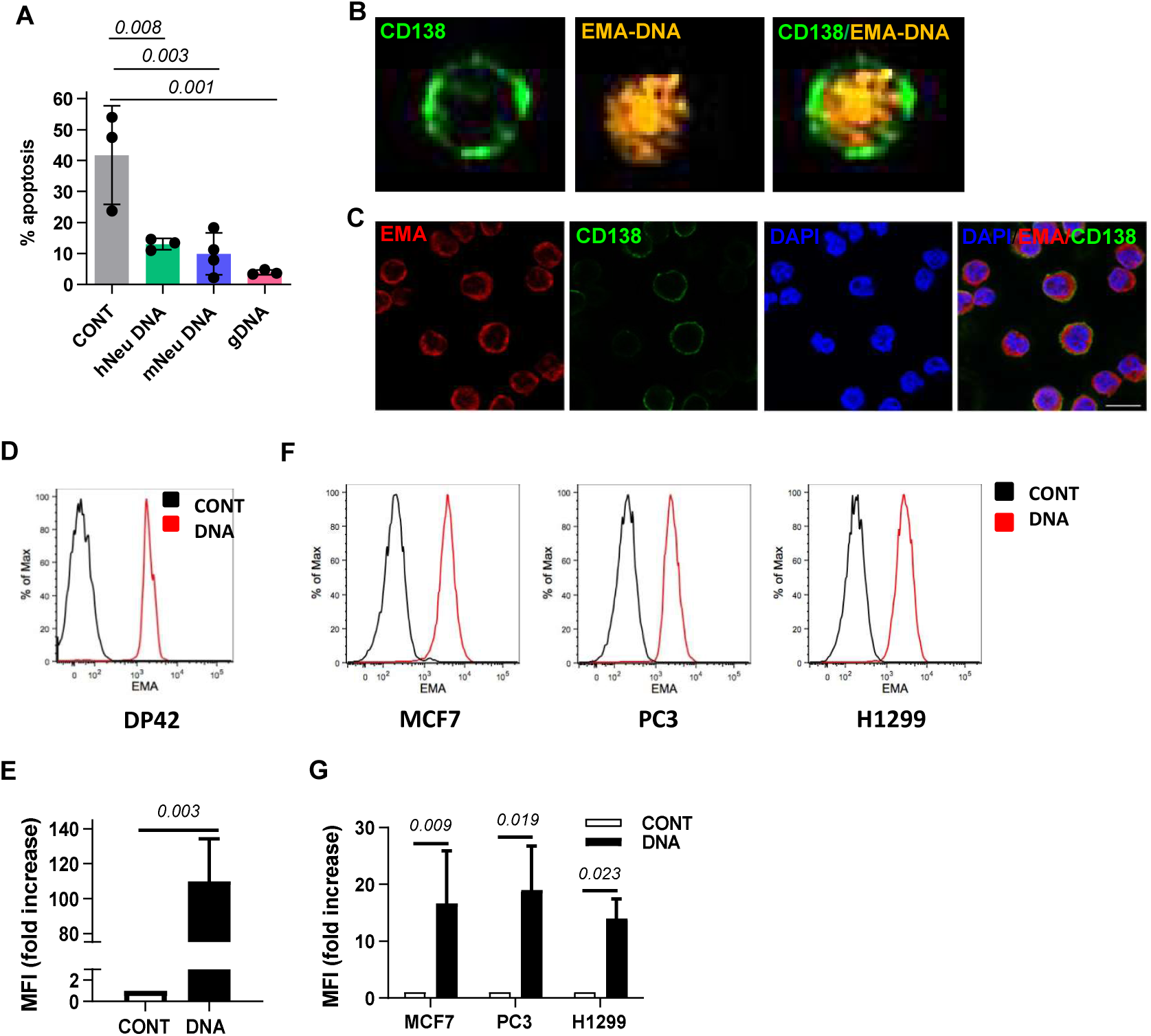
Uptake of cfDNA by tumor cells. (**A**) Doxorubicin-specific apoptosis of human H929 MM cells treated in the absence of DNA (CONTrol), in the presence of DNA isolated from supernatants collected from human or mouse neutrophil-MM cell cultures (hNeu DNA or mNeu DNA, respectively), or gDNA isolated from human neutrophils. Statistics: one-way ANOVA. (**B,C**) An ability of CD138^+^ MM cells to take up EMA-labeled DNA was evaluated by: (**B**) imaging flow cytometry (Amnis). Shown are representative images of one U266 cell. Data from at least 7,500 cells were acquired in each of three independent experiments; (**C**) confocal microscopy of MM1.S cells. Scale bar, 10 µm. (**D-G**) Uptake of EMA-labeled DNA by mouse DP42 cells (**D,E**) and indicated human cancer cell lines (**F,G**) after overnight incubation was evaluated by flow cytometry. Representative histograms (**D,F**) and geometric mean and SD values of at least three independent experiments (**E,G**) are shown. CONT – control (no cfDNA added). Statistics: paired two-tailed Student’s *t* test.

To investigate how cfDNA may interact with MM cells, DNA was labeled with ethidium monoazide bromide (EMA) and incubated with human MM cells. EMA-labeled DNA was internalized by human MM cells as detected by imaging flow cytometry (**Figure 4B**). Confocal microscopy demonstrated that this DNA localized preferentially in the cytoplasm of tumor cells (**Figure 4C**). DNA uptake was also observed in mouse MM cells (**Figure 4D,E**) and several human cancer cell lines including breast, prostate and lung cancer cells (**Figure 4F,G**) as evaluated by flow cytometry. Since MM-induced NETs contained gDNA and gDNA had a similar chemoprotective effect as cfDNA purified from neutrophil supernatants, we used gDNA isolated from mouse cells for all subsequent experiments (and refer to it as DNA).

Tumor cells began to take up DNA as soon as 1 hour after its addition and maintained this DNA inside the cell for at least 5 days (**Figure 5A**). Previous studies demonstrated that tumor-derived DNA delivered into dendritic cells (DCs) by transfection with lipofectamine activated STING signaling (Woo et al., 2014). In agreement with these data, we observed the presence of extracellular, EMA-labeled DNA in transfected DCs. However, without transfection, DCs were not able to take up DNA (**Figure S4A,B**) suggesting different mechanisms of DNA uptake by tumor cells and DCs. Tumor cells were able to take up 70 kDa Dextran, the commonly used marker of macropinocytosis. The amiloride derivative HMA and the P2 purinoreceptor antagonist suramin, each of which inhibits micropinocytosis, both blocked 70 kDa Dextran uptake (**Figure 5B**). We hypothesized that DNA entry into tumor cells is also mediated by endocytosis. Treatment of tumor cells with HMA or suramin almost completely prevented uptake of DNA (**Figure 5C**). Thus, MM cells are endowed with the ability to take up cfDNA via endocytosis.

**Figure 5.**
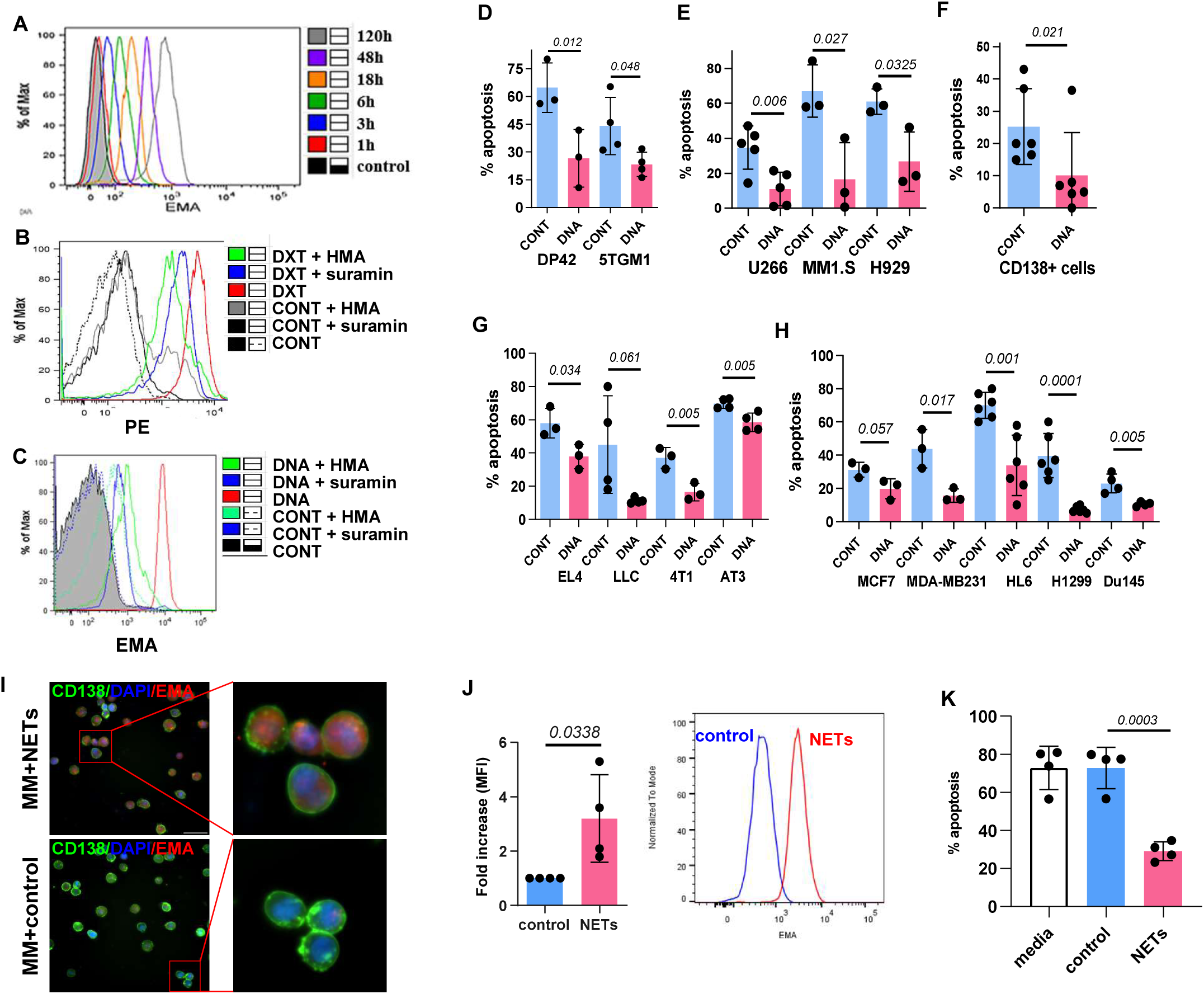
NETs and cfDNA protect tumor cells from apoptosis induced by doxorubicin. (**A**) Kinetics of EMA-labeled DNA (5 μg/mL) uptake by DP42 cells was evaluated by flow cytometry. (**B,C**) An effect of pharmacological inhibition of endocytosis on uptake of 70 kDa Dextran-TRITC (**B**) or EMA-labeled DNA (**C**) by human MM cells. Typical histograms are shown. Experiment was repeated at least three times with similar results. CONTrol – no cfDNA added. (**D-H**) Doxorubicin-induced apoptosis of mouse (**D**) or human (**E**) MM cell lines, primary human MM cells isolated from BM of 6 MM patients (**F**) or mouse (**G**) and human (**H**) cancer cell lines cultured without (CONTrol) or with cfDNA. Individual values, mean and SD values combined from 3 independent experiments are shown. Statistics: two-tailed Student’s *t* test. (**I-K**) NETs or control neutrophil supernatant (control) were collected from PMA-stimulated or unstimulated human neutrophils, respectively, and added to MM cells. An ability of H929 cells labeled with anti-CD138 FITC-conjugated antibodies to take up EMA-labeled NETs was evaluated by (**I**) microscopy; magnification, 10x; scale bar, 100 µm, or (**J**) flow cytometry. Representative graph (left) and fold increase in MFI between MM cells treated with NETs over control (right) are shown. (**K**) Doxorubicin-induced apoptosis of H929 cells treated in a presence of NETs, control neutrophil supernatant, or complete media was evaluated. Individual values, mean and SD values combined from 4 independent experiments are shown. Statistics: two-tailed Student’s *t* test.

The presence of DNA significantly reduced the cytotoxic effect of doxorubicin in mouse (**Figure 5D**) and human (**Figure 5E**) MM cell lines, and human CD138^+^ primary MM cells (**Figure 5F**). The chemoprotective effect of DNA was not limited to doxorubicin, as a similar effect was observed with other anthracycline antibiotics, including daunorubicin, epirubicin, and idarubicin (**Figure S5A**). Addition of DNA also protected different mouse (**Figure 5G**) and human (**Figure 5H**) lung cancer, breast cancer, lymphoma, and leukemia cells from doxorubicin-induced apoptosis. Notably, addition of cfDNA derived from apoptotic cells did not protect tumor cells from doxorubicin (**Figure S5B,C**).

We investigated whether tumor cells can take up NETs. We collected NETs from PMA-stimulated neutrophils and control supernatant from unstimulated neutrophils (**Figure S5D,E)**. Uptake of EMA-labeled NETs by MM cells was observed by microscopy (**Figure 5I**) and flow cytometry (**Figure 5J**). The presence of purified NETs protected human tumor cells from doxorubicin-induced apoptosis (**Figure 5K**).

gDNA and purified mtDNA similarly protected MM cells from doxorubicin-induced apoptosis (**Figure S5F**), whereas the addition of RNA did not reduce MM cell chemosensitivity (**Figure S5G**). The chemoprotective effect of DNA was not mediated through unmethylated CpG DNA motifs recognized by Toll-like receptor 9 (TLR9), as the presence of stimulatory CpG oligodeoxynucleotides (ODN) or control CpG-ODN equally protected human MM cells from doxorubicin (**Figure S5H**). TLR9 antagonist ODN did not prevent the chemoprotective effect of DNA (**Figure S5I**). Both Cy5-labeled ssDNA ODN and ssRNA oligonucleotides were internalized by MM cells and localized in the cytoplasm (**Figure S6A,B**). However, in contrast to DNA and ssDNA ODN, the presence of ssRNA oligonucleotides did not affect the chemosensitivity of MM cells (**Figure S6C**). No co-localization of ssDNA ODN with the lysosomal protein LAMP1 was found in MM cells (**Figure S6D**).

We investigated the possibility that upon entering tumor cells cfDNA activates signaling that protects tumor cells from doxorubicin-induced apoptosis. For that, we evaluated the expression of several apoptosis related proteins in MM cells treated with DNA. No effect on the level of BCL2, BCL-xL, MCL1, and survivin was found (**Figure S7A**). No differences were also observed in MAP kinases phosphorylation (**Figure S7B**).

Since anthracyclines are intercalating agents, we tested the possibility that they could bind NET-derived DNA. Doxorubicin is naturally fluorescent and therefore can be easily visualized using fluorescent microscopy. Indeed, addition of doxorubicin to neutrophils undergoing NETosis demonstrated its co-localization with NETs (**Figure 6A**). No binding to NETs was observed with paclitaxel, a chemotherapeutic that interacts with microtubules but not DNA (**Figure 6B**). In agreement with these data, addition of cfDNA did not protect human MM cells from paclitaxel or the proteasome inhibitor bortezomib (**Figure 6C**).

**Figure 6.**
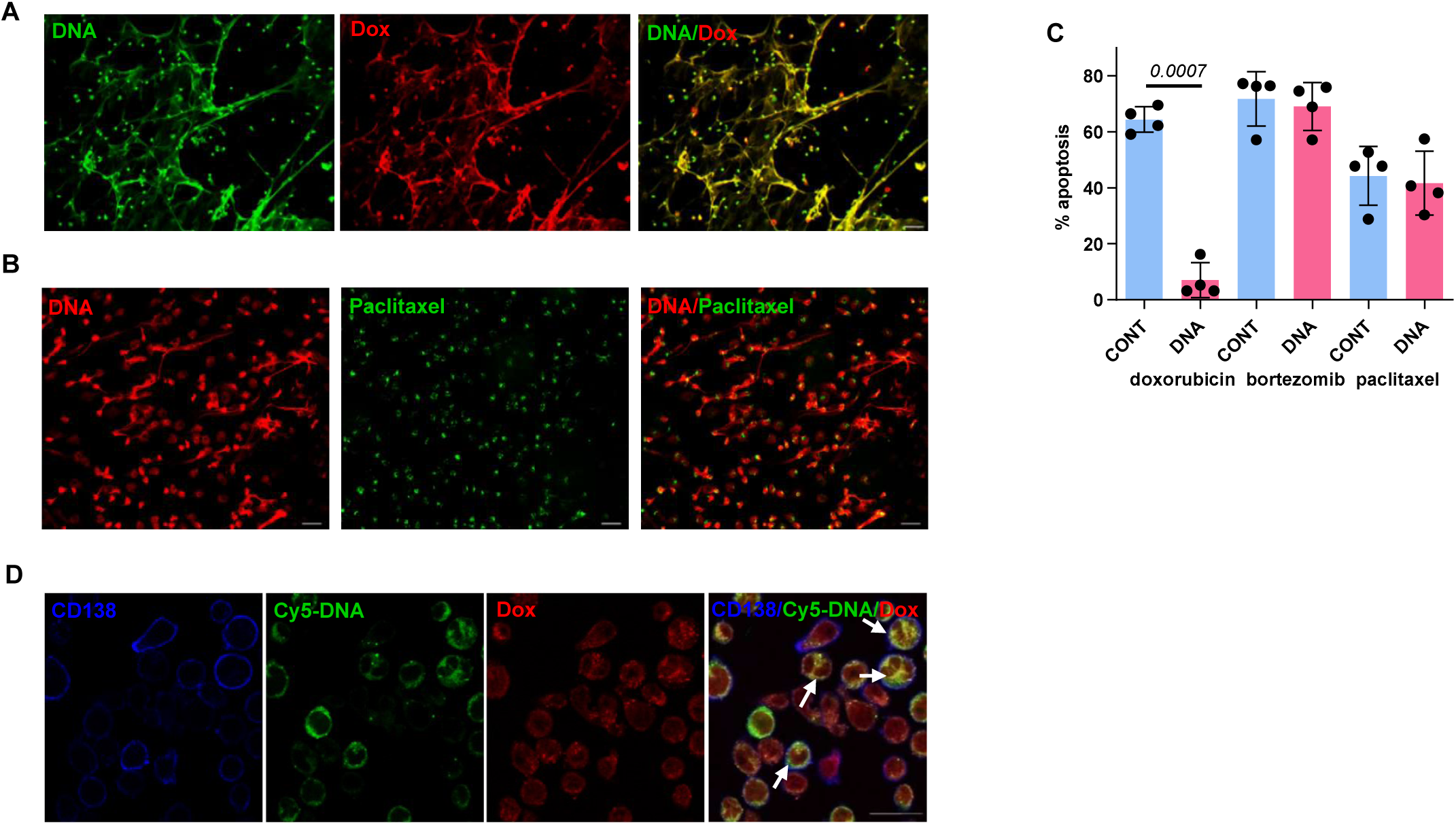
NETs and cfDNA bind doxorubicin. (**A**) An ability of doxorubicin (Dox) to bind NET DNA as evaluated by fluorescent microscopy. PMA-stimulated human neutrophils were treated for 1h with 1 µM Dox followed by staining with SYTOX Green Nucleic Acid stain (SYTOX). Representative images from one of four independent experiments. Magnification, 20x, Scale bar, 100 µm. (**B,C**) DNA does not protect tumor cells from paclitaxel. (**B**) Representative images showing localization of Oregon Green 488-conjugated Paclitaxel after 1h treatment of PMA-stimulated human neutrophils stained with SYTOX Orange Nucleic Acid Stain (SYTOX). Magnification 20x; scale bar, 50 µm. Images were acquired using upright microscope. (**C**) Chemotherapy-induced apoptosis of H929 cells treated in the presence (DNA) or absence (CONTrol) of cfDNA. Individual results, mean and SD values are shown. Statistics: two-tailed paired Student’s *t* test. (**D**) Intracellular binding of Dox with Cy5-DNA ODN was evaluated by confocal microscopy in MM1.S cells labeled with anti-CD138 antibody. Representative images from one of three independent experiments. Scale bar, 50 µm. Arrows indicate co-localization of Dox with DNA staining.

Doxorubicin is typically quickly transported to the nucleus upon entering cells. However, internalized cfDNA was able to entrap doxorubicin in the cytoplasm of tumor cells (**Figure 6D**), thus limiting its nuclear availability and reducing its cytotoxic effect.

### Therapeutic targeting NETs and cfDNA enhance the anti-tumor effect of doxorubicin

To confirm that neutrophil-derived cfDNA is indeed a factor responsible for the reduced sensitivity of MM cells to doxorubicin, we treated MM cells with doxorubicin in the presence of deoxyribonuclease I (DNase I). Addition of DNase I reversed the protective effect of mouse neutrophils on doxorubicin-induced apoptosis of MM cells (**Figure 7A**). In human cells, DNase I significantly increased the cytotoxicity of doxorubicin in MM cells cultured with BM plasma from relapsed MM patients, containing large amount of DNA (**Figure 7B**). To investigate whether targeting cfDNA would enhance the effect of doxorubicin *in vivo*, we used the recombinant DNase I Pulmozyme in a syngeneic mouse model of MM. As expected, administration of doxorubicin delayed MM progression. Treatment with Pulmozyme alone did not demonstrate an anti-tumor effect. However, the mice that received doxorubicin in combination with Pulmozyme never developed MM symptoms (**Figure 7C, D**). The PAD4 inhibitor BMS-P5 (Li et al., 2020) was further used to evaluate the potential of targeting NETs as a strategy for enhancing the effect of doxorubicin. Monotherapy with BMS-P5 or doxorubicin demonstrated a moderate anti-tumor effect in a syngeneic MM model. However, the combination of these two agents resulted in potent anti-tumor activity evident by delayed onset of paralysis and prolonged survival (**Figure 7E, F**). Taken together, these results suggest that targeting cfDNA or NETs would significantly improve the anti-tumor effect of anthracyclines.

**Figure 7.**
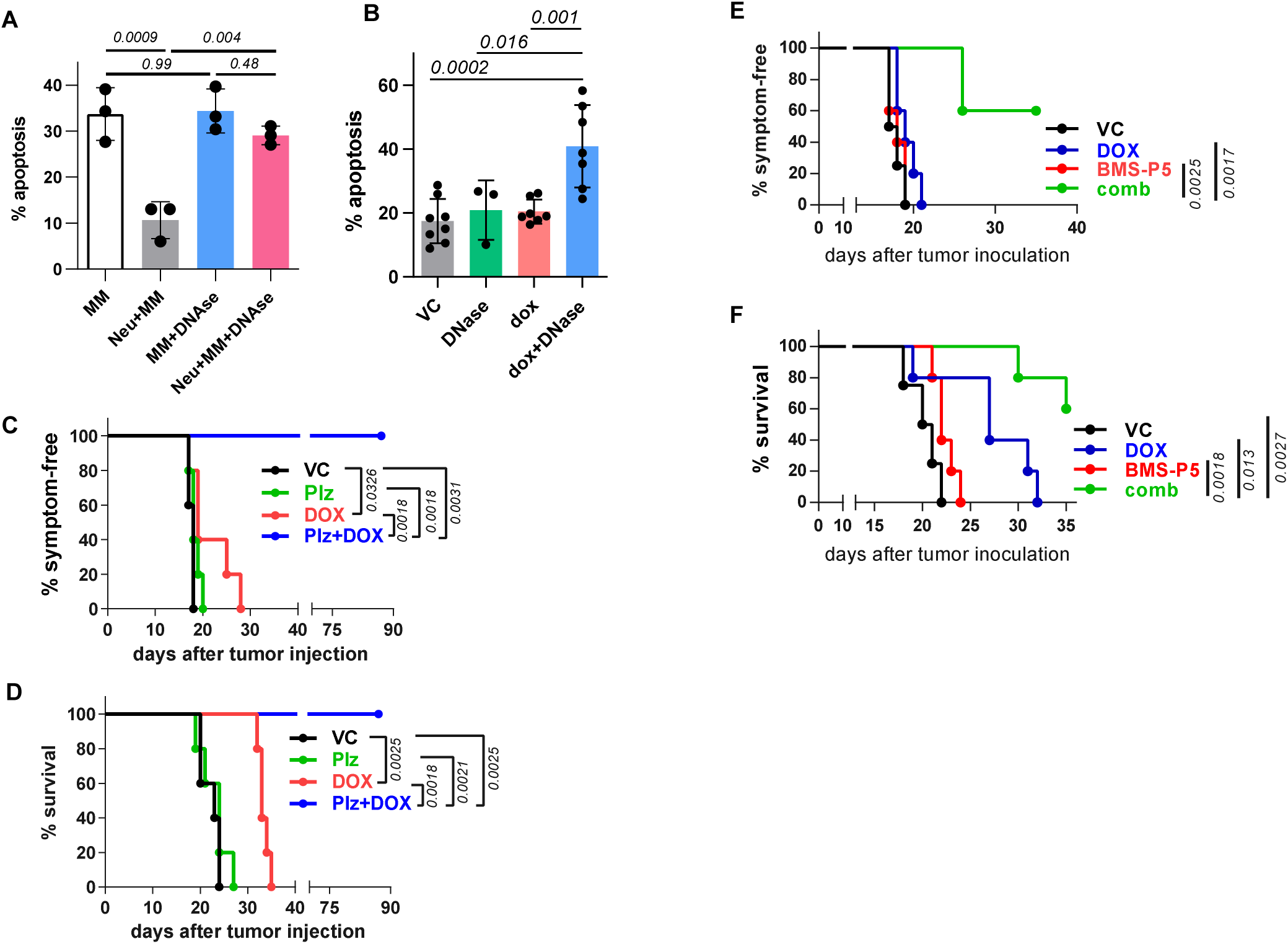
Neutrophils protect MM cells by releasing NETs. (**A**) Doxorubicin-induced apoptosis of mouse DP42 MM cells cultured in the presence or absence of BM neutrophils with or without addition of 500 U/mL DNase I. (**B**) Apoptosis of human MM cells kept in BM plasma from MM patients and incubated overnight in the presence or absence of DNase I followed by 24h treatment with dox or vehicle control (vc, PBS). Statistics: one-way ANOVA. (**C,D**) Onset of symptoms (**C**) and survival (**D**) of tumor-bearing DP42 mice (n=5 per group) treated with vehicle control (VC), doxorubicin (DOX), Pulmozyme (Plz), or combination of Pulmozyme and doxorubicin (Plz+DOX). Statistics: log-rank test. (**E,F**) Onset of symptoms (**E**) and survival (**F**) of tumor-bearing DP42 mice (n=5 per group) treated with VC, DOX, PAD4 inhibitor BMS-P5, or combination of BMS-P5 and doxorubicin. Statistics: log-rank test.

## Discussion

Our study has demonstrated a novel mechanism by which neutrophils in the BM microenvironment protect tumor cells from anthracycline chemotherapy agents.

CD11b^+^Ly6C^lo^Ly6G^+^ neutrophils are the most abundant cells in the BM. We found that neutrophils from TB mice produced significantly more NETs than neutrophils from TF mice. These data are in agreement with a recent study suggesting that circulating neutrophils isolated from TB mice and stimulated *in vitro* with platelet-activating factor are more prone to NET formation compared to neutrophils isolated from wild-type mice (Demers et al., 2012). While in TF mice CD11b^+^Ly6C^lo^Ly6G^+^ cells are represented by neutrophils, in TB mice cells with this phenotype are comprised largely of pathologically activated PMN-MDSC (Veglia et al., 2021). We and others have previously reported that growth of MM in mice and humans is accompanied by accumulation of PMN-MDSC in the BM (Ramachandran et al., 2016). In mice, PMN-MDSC share the CD11b^+^Ly6C^lo^Ly6G^+^ phenotype with neutrophils, but exhibit distinct functional and biochemical characteristics, and have an ability to suppress immune responses (Condamine et al., 2016; Gabrilovich and Nagaraj, 2009). It is possible that PMN-MDSC have a stronger ability to form NETs compared to neutrophils. We have recently reported that neutrophils could phenotypically be distinguished from PMN-MDSC in the tissues of TB mice based on the expression of CD14 (Veglia et al., 2021). This information would allow a comparison of the NET-forming ability of neutrophils and PMN-MDSC. Similar to the mouse system, we found that BM neutrophils from patients with MM formed substantially more NETs in response to stimulation with MM cells than neutrophils from the BM of healthy donors.

Increased levels of circulating cfDNA in PB of cancer patients has been described and extensively explored as a tool for cancer screening and as a prognostic biomarker (Bronkhorst et al., 2019). However, a significant proportion of PB cfDNA consists of non-mutated DNA. Thus, some studies demonstrated that tumor-derived cfDNA (circulating tumor DNA) may constitute only a small fraction (less than 1%) of total cfDNA, suggesting a different source for most of this DNA (Diaz and Bardelli, 2014). NETosis is a unique mechanism that may be responsible for the presence of a significant amount cfDNA in the PB of cancer patients. Citrullination of histone H3 is a hallmark of NETosis. Consistently with other reports, we demonstrated co-localization of citrullinated histone H3 with DNA in NETs. In addition, we found that not only levels of extracellular DNA but also levels of DNA bound to citrullinated histone H3 were significantly increased in the BM of TB hosts.

We demonstrated that uptake of extracellular DNA protected different types of tumor cells from anthracyclines. Clathrin-dependent, caveolae-dependent, and macropinocytosis are the main endocytic mechanisms. Our data showed that uptake of cell-free DNA by tumor cells relies on macropinocytosis, as pharmacological inhibition of this pathway significantly blocked 70 kDA Dextran-TRITC and extracellular DNA entry. Immature DCs are characterized by robust endocytic activity. However, in contrast to tumor cells, DCs were not able to take up extracellular DNA unless it was delivered by transfection. These results are in agreement with a study by Woo et al, which reported that tumor cell-derived DNA delivered into DCs by transfection initiated DNA sensing and a type I interferon response (Woo et al., 2014). DCs do not express caveolin-1 and 2 (Carter et al., 2011), the main structural proteins of caveolae. It is possible that the lack of caveolins in DCs may explain the inability of these cells to take up extracellular DNA. Recently, de Mingo Pulido and co-authors demonstrated that expression of TIM3 may regulate uptake of extracellular DNA by cDC1 (de Mingo Pulido et al., 2021). Thus, tumor DNA was taken up by DC1 poorly. However, the uptake was dramatically upregulated upon blocking TIM3 (de Mingo Pulido et al., 2021). TIM3 is highly expressed by MM cells (Liu et al., 2020). Interestingly, high TIM3 expression did not prevent uptake of extracellular DNA by tumor cells.

Our experiments demonstrated that the effect of DNA on protection of tumor cells from anthracyclines was most likely mediated not by specific signaling, but rather by sequestration of the drug in the cytoplasm that prevented it from reaching the nucleus.

NETs are not the only source of cfDNA. CfDNA could also originate from cells undergoing other mechanisms of cell death such as apoptosis or necrosis. However, our data demonstrated that cfDNA derived from apoptotic cells did not protect tumor cells from anthracyclines. This could be explained by the fact that the size of DNA fragments following apoptotic cell death are short due to the digestions by DNase. Inability to detect apoptotic DNA using agarose gel confirms that. A similar effect was observed when genomic DNA was digested with DNase I prior to its addition to the tumor cells. In blood, cfDNA is immediately digested by DNase I. It is possible that NET DNA is less accessible to DNase I due to its complexes with protein, and therefore it may not undergo rapid degradation. Increasing DNase I concentration by administration of the enzyme to mice overcame this limitation and significantly improved anti-tumor response to doxorubicin treatment.

Our experimental data are primarily from myeloma cells, and anthracyclines are no longer commonly used in the initial lines of therapy for patients with multiple myeloma. However, doxorubicin is commonly used in multiagent chemotherapy regimens for aggressive relapsed multiple myeloma (particularly in the D-PACE regimen) (Lakshman et al., 2018). Improvements in the therapeutic efficacy of doxorubicin could increase utilization of this chemotherapy agent in the therapy of relapsed and refractory myeloma and improve patient outcomes.

In summary, we have demonstrated a new mechanism by which the microenvironment reduces the sensitivity of tumor cells to chemotherapeutics. Targeting this mechanism could be beneficial for treatment of cancer patients.

## Acknowledgments

Support for this work was provided by NIH/NCI grant R01CA196788 (to Y.N.) and T32CA009171 (to S.E.H.). Support for the Shared Resources utilized in this study was provided by Cancer Center Support Grant P30CA010815 to The Wistar Institute. We would also like to thank Dr. Qin Liu for assistance with data analysis, Frederick Keeney for assistance with microscopy, and Dr. Rachel E. Locke for helping with the preparation of the manuscript.

## Author contributions

CL and SEH performed experiments, analyzed the data, and prepared the draft of the manuscript. ML, HD, and RK performed experiments. MR provided technical assistance. LB coordinated the acquisition of clinical samples. DTV selected patients, provided clinical samples and analyzed clinical data. DVT and DG discussed the results and reviewed the manuscript. YN was responsible for project design, overall supervision, evaluation of experimental data, and writing the manuscript.

## Conflict of interest statement

The authors declare no competing interests.

## Methods

### Human subjects and samples

Collection of samples from patients with MM and donors at the Abramson Cancer Center at the University of Pennsylvania was approved by the Institutional Review Board (IRB) of the University of Pennsylvania. All patients signed IRB-approved consent forms. Samples were collected from 20 patients with newly diagnosed untreated MM and 40 patients with relapsed/refractory MM. This cohort included 20 females and 40 males, 34-83 years of age.

### Isolation of human primary cells

Neutrophils were isolated from BM samples using Percoll (GE Healthcare) gradient from cells pelleted following Ficoll-Paque (GE Healthcare) gradient centrifugation. Neutrophils were isolated from PB using Histopaque (Sigma-Aldrich) gradient centrifugation. Primary MM cells were isolated from BM mononuclear fraction by positive selection of CD138 cells using CD138 MicroBeads and MACS columns (all from Miltenyi Biotec).

### Tumor cell lines and reagents

Tumor cell lines: DP42 (mouse multiple myeloma), 5TGM1 (mouse multiple myeloma), AT3 (mouse breast cancer) were kindly provided by Dr. Brian Van Ness, Dr. Lori Hazlehurst, and Dr. Barry Hudson, respectively. KMS-11 and KMS-28BM (both human multiple myeloma) cells were obtained from JCRB. U266, MM1.S, H929 (NCI-H929), RPMI-8226 (all human multiple myeloma), MCF7 and MDA-MB231 (human breast cancer), HL60 (human leukemia), H1299 (human non-small cell lung cancer), Du145 (human prostate cancer), EL4 (mouse lymphoma), 4T1 (mouse breast cancer), and LLC (mouse Lewis carcinoma) were from ATCC. MM cells were cultured in RPMI-1640 and solid cancer cells in DMEM medium; all supplemented with 10% fetal bovine serum (FBS, Atlanta Biologicals) and penicillin/streptomycin (Gibco). DP42 cells were also supplemented with 0.5 ng/mL murine IL-6 (Biolegend). All cell lines were periodically tested for mycoplasma contamination by Universal Mycoplasma Detection kit (ATCC). Reagents and antibodies information is provided in **Table 1**.

**Table 1.** Reagents and antibodies used in the study.

| **Company** | **Reagents/Antibody name** | **Catalog number** |
| --- | --- | --- |
| Sigma | doxorubicin | D1515 |
|  | DNase I | D5025 |
|  | PMA | P1585 |
|  | 5-(N,N-Hexamethylene)amiloride (HMA) | A9561 |
| Sellekchem | daunorubicin | S3035 |
|  | epirubicin | S1223 |
|  | idarubicin | S1228 |
| Miltenyi Biotech | anti-Ly6G biotin, mouse | 130-101-884 |
|  | Streptavidin Microbeads | 130-048-101 |
|  | CD138 Microbeads, human | 130-051-301 |
| BD Biosciences | APC Rat Anti-Mouse Ly6G | 560599 |
|  | PE Rat Anti-Mouse Ly6G | 551461 |
| | FITC Mouse IgG1, $\kappa$ isotype, control | 555909 |
|  | FITC mouse anti-human CD138 | 552723 |
|  | purified mouse anti-human CD107a (LAMP1) | 555798 |
| BioLegend | Brilliant Violet 421 anti-human CD138 | 356515 |
|  | APC Annexin V | 640941 |
| | PE Mouse IgG2b, $\kappa$ isotype, control | 400314 |
|  | Brilliant Violet 421 anti-mouse/human CD11b | 101236 |
|  | PE/Cy7 anti-mouse CD38 | 102717 |
| Cayman Chemical | Suramin (sodium salt) | 11126 |
| R&D Systems | recombinant mouse IL-6 protein | 406-ML |
|  | recombinant mouse GM-CSF protein | 415-ML |
|  | recombinant mouse Syndecan-1 Protein, CF | 3190-SD-050 |
| Invitrogen | myeloperoxidase polyclonal antibody (MPO) | PA5-16672 |
| ThermoFisher Scientific | Sytox Green Nucleic Acid Stain | S7020 |
|  | Sytox Orange Nucleic Acid Stain | S11368 |
|  | POPO-1 iodide | P3580 |
|  | ethidium monoazide bromide (EMA) | P22310 |
|  | paclitaxel, Oregon Green 488 conjugate | P22310 |
|  | DAPI | D3571 |
|  | Dextran, Tetramethylrhodamine, 70,000 MW, Lysine Fixable | D1818 |
| Qiagen | QIAquick PCR Purification Kit | 28104 |
| Abcam | anti-Histone H3 (citrulline R2+R8+R17) antibody | ab5103 |
|  | mitochondrial DNA isolation kit | 65321 |
|  | GM 6001 | ab120845 |
| InvivoGen | TLR9 antagonist ODN TTAGGG | Tlrl-ttag151 |
|  | human TLR9 agonist ODN 2395, ODN 2006, ODN 2336 | tlrl-2395, tlrl-2006, tlrl-2336 |
|  | ODN 2395 control, ODN 2006 control, ODN 2336 control | tlrl-2395c, tlrl-2006c, tlrl-2336c |
| Applied Biosystems | High Capacity cDNA Reverse Transcription Kit | 4368814 |
|  | Power SYBR green PCR Master Mix | 4367659 |
| Omega Bio-Tek | E.Z.N.A. Total RNA Kit I | R6834-02 |
| Santa Cruz | β-Actin Antibody (C4) | sc-47778 |
| Cell Signaling Technology | Bcl-xL (54H6) Rabbit mAb | 2764T |
|  | Bcl-2 (D17C4) Rabbit mAb | 3498T |
|  | Survivin (71G4B7) Rabbit mAb | 2808T |
|  | Mcl-1 (D2W9E) Rabbit mAb | 94296S |
|  | SAPK/JNK Antibody | 9252T |
|  | Phospho-SAPK/JNK (Thr183/Tyr185) (81E11) Rabbit mAb | 4668T |
|  | p38 MAPK (D13E1) Rabbit mAb | 8690S |
|  | Phospho-p38 MAPK (Thr180/Tyr182) (D3F9) Rabbit mAb | 4511S |
|  | Akt (pan) (11E7) Rabbit mAb | 4685S |
|  | Phospho-Akt (Ser473) (D9E) Rabbit mAb | 4060T |
|  | p44/42 MAPK (Erk1/2) Rabbit mAb | 4695S |
|  | Phospho-p44/42 MAPK (Erk1/2) Rabbit mAb | 4370S |
| Tecan | anti-HMGB1 [DPH1.1] monoclonal antibody | REHM901 |

### Mouse models and treatment

All animal experiments were carried out in accordance with institutional guidelines and were approved by the Institutional Animal Care and Use Committee of the Wistar Institute. C57BL/6 and FVB/N mice were purchased from Charles River Laboratories and were crossed to obtain F1 progeny of mixed C57BL/6×FVB/N background. In these mice, MM tumors were established by i.v. inoculation of DP42 cells (5×10^3^ in a volume of 100 µl of PBS) into the tail vein. 5TGM1 tumors were established in C57BL/KaLwRij mice by i.v. inoculation of 5TGM1 cells (1×10^6^) into the tail vein. Six to ten weeks old male and female mice were used for experiments. TLR4 KO mice were purchased from the Jackson Laboratory and RAGE KO mice were provided by Dr. John Hoidal (University of Utah).

For *in vivo* treatment studies, DP42-bearing mice were split into groups and treated with Doxorubicin (Pfizer), Pulmozyme (Genentech, San Francisco, CA), PAD4 inhibitor BMS-P5 (kindly provided by BMS, NJ), or combination of Doxorubicin with either Pulmozyme or BMS-P5. Control group of mice received PBS. Doxorubicin was given at a dose of 3 mg/kg, i.v., every 4 days beginning on day 10 after tumor cell injection. Mice received a total of 5 Doxorubicin injections. BMS-P5 was given at a dose of 50 mg/kg, via oral gavage, twice a day, daily, beginning on day 5 after tumor cell injection. Pulmozyme was administered at a dose of 5 mg/kg, i.v., 15 min prior to each Doxorubicin injection and at a dose of 2.5 mg/kg, i.p., 4h, 8h, and overnight following each Doxorubicin injection. Onset of symptoms (paralysis and hunched posture) was monitored. Survival of mice was determined as they were euthanized at the humane endpoint.

### Isolation of mouse neutrophils

Neutrophils were isolated from BM cells using MACS technique. Cells were labeled with biotin-Ly6G antibody followed by incubation with Streptavidin-conjugated MicroBeads and separation on MACS columns (Miltenyi Biotech). The purity of the isolated Ly6G population was more than 98% as evaluated by flow cytometry.

### Isolation and labeling of DNA

ExtDNA was purified from supernatants collected from MM-neutrophil co-cultures; gDNA was isolated from BM neutrophils using standard phenol-chloroform extraction. Mitochondrial DNA was isolated from HL60 cells using Mitochondrial DNA isolation kit.

For labeling, EMA was added to DNA in a final concentration of 5 µg/mL and exposed to light for 20 min. DNA was then washed with ethanol to ensure removal of non-bound EMA.

### NET detection

Neutrophils cultured in 24 flat-bottom plates with or without MM cells placed in transwell inserts were fixed with 4% paraformaldehyde followed by staining with Sytox Green Nucleic Acid Stain at a final concentration of 250 nM. A Nikon Te300 inverted microscope equipped with a motorized XY stage was used to image NETs. Nikon Nis-Elements Ar software was used for image acquisition and was set up to acquire 25 random locations per well. The z-stacks per location were then combined into an extended depth focused image. The total NET area was calculated by segmenting each image using a defined threshold pixel intensity setting. The spot detection tool in NIS-Elements Ar was used to count the number of cells per field. The sum of the total NET area in the 25 random fields of view was divided by the total number of cells in the 25 fields of view to obtain NET area (µM/cell).

### Cell treatment

Human tumor cells were plated at a density of 0.5×10^6^ cells/mL and mouse tumor cells at a density of 0.5-1×10^6^ cells/mL. Cells were cultured overnight in the presence or absence of different types of extracellular DNA (all at concentrations of 5-10 µg/mL) followed by 24h treatment with chemotherapeutics and evaluation of apoptosis. DP42, 5TGM1, EL4, and LLC cells were treated with 1-2 µM doxorubicin. Human MM cells were treated with 1-2 µM doxorubicin or 2 µM daunorubicin, idarubicin, or epirubicin; MCF7 and MDA-MB231 cells were treated with 7.5-10 µM doxorubicin. HL-60, H1299, or Du145 cells were cultured overnight with or without gDNA and treated for 48h with 1-4 µM doxorubicin.

### Generation of mouse DCs

Mouse BM cells were plated at a density of 3×10^5^ cell/mL/well in RPMI-1640 medium supplemented with 10% FBS, penicillin-streptomycin, 50 µM 2-mercaptoethanol, 10 ng/mL GM-CSF, and 100 ng/mL Flt3L. Media and cytokines were replaced every 3 days. DCs were collected on day 12 of culture.

### Cell transfection

DCs and MM cells were plated at a density of 1×10^6^ cell/mL in RPMI-1640 medium supplemented with 10% FBS and 1% Antibiotic-Antimycotic. Cells were transfected with EMA-labeled DNA (1µg/well) using Lipofectamine 3000 (Invitrogen) for 18h according to manufacturer’s instructions.

### Quantification of cfDNA

Quantification of cfDNA was performed using Sytox^®^ Green Nucleic Acid Stain (ThermoFisher Scientific). Briefly, 20 µL of cell supernatants or plasma were added per well of 96 well plate followed by addition of Sytox^®^ Green Nucleic Acid Stain at a final concentration of 1 µM. Fluorescence of Sytox Green was quantitated using a Wallac Victor2 Microplate reader (Perkin Elmer).

### NET isolation

Human neutrophils were isolated and plated at a density of 2×10^6^cells/mL in phenol-free RPMI1640 supplemented with 2% FBS. Neutrophils were stimulated with 200 nM PMA for 4hrs then collected and spun at 450 x g at 4° for 5 min. Supernatant was collected and centrifuged at 18,000 x g at 4° for 15min. Supernatant was then discarded, and the remaining NETs were resuspended in PBS.

### Flow cytometry

Apoptosis of MM cells was evaluated using Annexin V binding assay. Briefly, cells were washed twice with ice cold PBS and once with binding buffer followed by staining with Annexin V-APC (BioLegend) and DAPI (Life Technologies). For surface staining, cells were labeled with specific antibodies for 20 min at 4°C in dark, washed with ice cold PBS, and resuspended in PBS containing DAPI. For Dextran or DNA uptake, human MM cells were preincubated with 50 µM HMA or 500 µM suramin for 30 min followed by addition of 10 µg/mL70 kDa Dextran conjugated with Tetramethylrhodamine (TRITC) or EMA-labeled DNA. Cells were incubated for 6h at +37°C, washed, and resuspended in PBS containing DAPI. At least 10,000 events were acquired using LSR II flow cytometer (BD). Data were analyzed using FlowJo software (TreeStar).

### Imaging flow cytometry

EMA-labeled DNA was added to MM cells and incubated overnight followed by staining with anti-human CD138-FITC antibody (BD, cat no. 552723). Presence of DNA was determined using an ImageStream^X^ Mark II instrument (Amnis Corporation, Seattle, WA). Data were acquired at 40× magnification utilizing Extended Depth of Field using INSPIRE acquisition software. Data from a minimum of 7,500 cells were collected for each sample and analyzed using IDEAS 6.0 software.

### Confocal Microscopy

MM1.S cells (0.5×10^6^ cells/mL) were plated on poly-L-Lysine coated coverslips (Fischer Scientific) and placed in a 24-well plate. Cy5-labeled DNA ODN (5’CY5-TTG CCT GCC ACT ACA ACA GTA TCT A) or Cy5-labeled RNA oligonucleotide (5’CY5-UUG CCU GCC ACU ACA ACA GUA UCU A) was added at a concentration of 10 µg/mL and left overnight at 37°C. Doxorubicin (1 µM) was added for 1h followed by fixation with 4% paraformaldehyde for 15 min at room temperature. Coverslips were washed with PBS and stained with CD138-BV421 (BioLegend, cat no. 356515, 1:200 dilution) for 20 min at room temperature. Coverslips were washed again, mounted onto slides using ProLong Gold antifade reagent (Life Technologies) and allowed to air dry before analysis.

In some experiments, MM1.S cells incubated overnight in the presence of Cy5-labeled DNA ODN were fixed with 4% paraformaldehyde and then labeled with anti-LAMP1 antibody (BD, cat no. 555798, 1:100 dilution) followed by staining with 1 µg/mL DAPI.

To evaluate localization of extracellular DNA inside MM cells, EMA-labeled DNA was added to MM1.S cells at a final concentration of 5 µg/mL and incubated overnight. Cells were then labeled with CD138-FITC antibody (BD, cat no. 552723) and fixed with 4% paraformaldehyde followed by staining with 1 µg/mL DAPI.

Images were acquired using a Leica TCS SP5 II scanning laser confocal microscope.

### Microscopy

BM neutrophils kept with or without DP42 cells placed in TW insert were blocked with 5% BSA for 30 min and stained with antibody against citrullinated Histone 3 for 2h at room temperature. Cells were labeled with Alexa-647-conjugated anti-rabbit antibody and fixed with 4% paraformaldehyde followed by staining with POPO-1 Nucleic Acid Stain. Images were acquired using Nikon 80i upright microscope.

### Western blotting

Western blotting was performed according to the standard protocol. MM cells were lysed in RIPA buffer supplemented with 1× Halt protease and phosphatase inhibitor (Cat. #1861281, ThermoFisher Scientific). Membranes were blocked in 5% milk for 1h at room temperature and then incubated overnight at +4°C with the primary antibodies against indicated proteins. Membranes were then washed, incubated with corresponding secondary antibodies, and developed using ECL (Bio-Rad, Hercules, CA).

### Real-time quantitative PCR

PCR was performed to determine a presence of male-specific sex-determining region of the Y chromosome (SRY). Genomic DNA was isolated from DP42 cells originally derived from the male mouse. ExtDNA was extracted from supernatants collected from co-culture of neutrophils from female mice with DP42 cells. Real-time PCR was performed on an ABI 7500 Fast instrument (Applied Biosystems) using 100 ng of the template and Power SYBR Green PCR Master Mix (Applied Biosystems). Expression of SRY was normalized to the expression of GAPDH. Proportion of Y chromosome DNA in supernatants from neutrophil-DP42 co-cultures was calculated. Primers used are listed in Supplementary Table 2.

### Conventional PCR

The presence of nuclear-and mt-specific genes was determined using primers listed in **Table 2**. The amplification parameters were as follows: 98°C for 10sec, 60.1°C for 30sec, 72°C for 1.5 min, 30 cycles. The PCR products were separated on 3% agarose gel and visualized using staining with ethidium bromide.

**Table 2.**
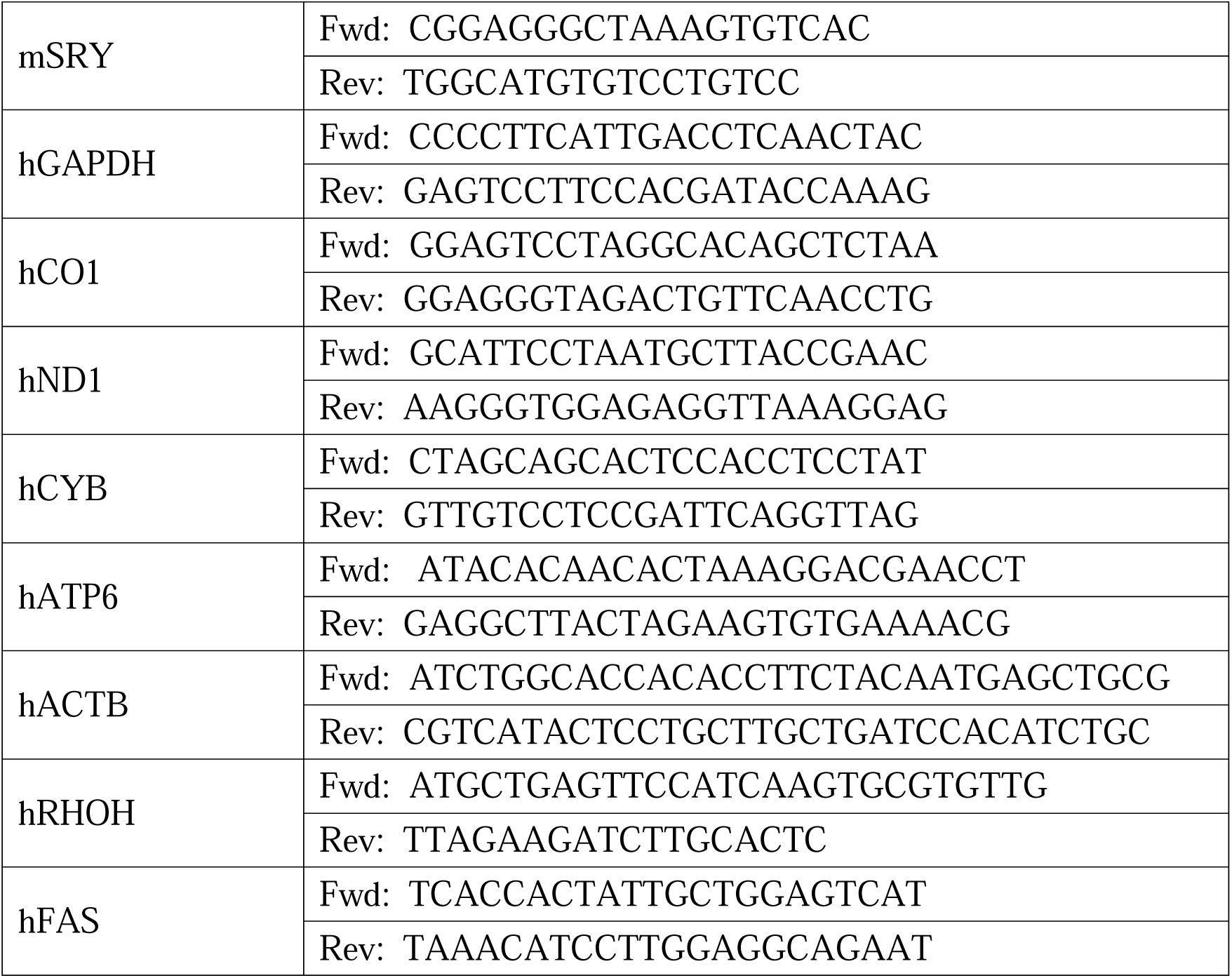
Primers used in the study.

### Statistics

Statistical analysis was performed using GraphPad Prism 5 software (GraphPad Software, Inc, La Jolla, CA). Student *t* test, one-way ANOVA, or Mann-Whitney test were used to determine differences between treatment groups. A log-rank test was used for the comparison of survival curves. A *p* value of less than 0.05 was considered statistically significant.

## Notes

### Competing Interest Statement

The authors have declared no competing interest.

