## Supplemental Figures for "Neutrophil extracellular traps promote tumor chemoresistance to anthracyclines"

**Figure S1**

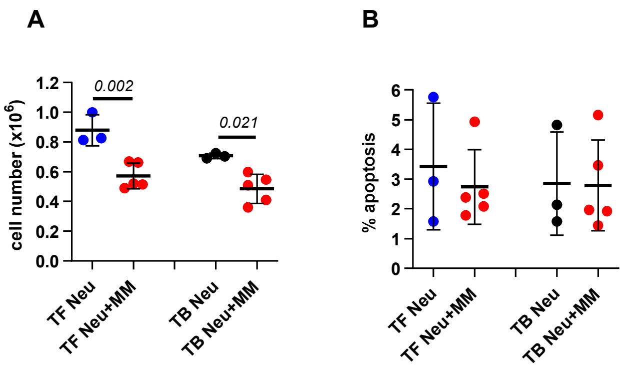


**Figure S1. MM cells do not induce apoptosis of neutrophils, Related to Figure 1.** Cell number evaluated by Trypan blue exclusion (A) and apoptosis evaluated by Annexin V binding assay (B) of neutrophils from TF and TB DP42 mice cultured either alone or in the presence of MM cells separated by Transwell insert Individual values, mean, and SD are shown. Statistics: one-way ANOVA.

**Figure S2**

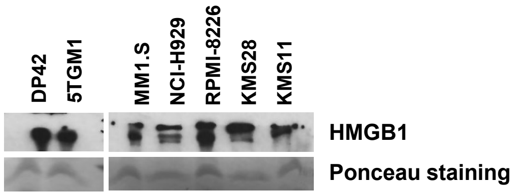


**Figure S2. Production of HMGB1 by MM cells, Related to Figure 2.** HMGB1 was detected in supernatants collected from indicated MM cells by western blotting. Ponceau staining of membrane was performed to confirm protein loading.

**Figure S3**

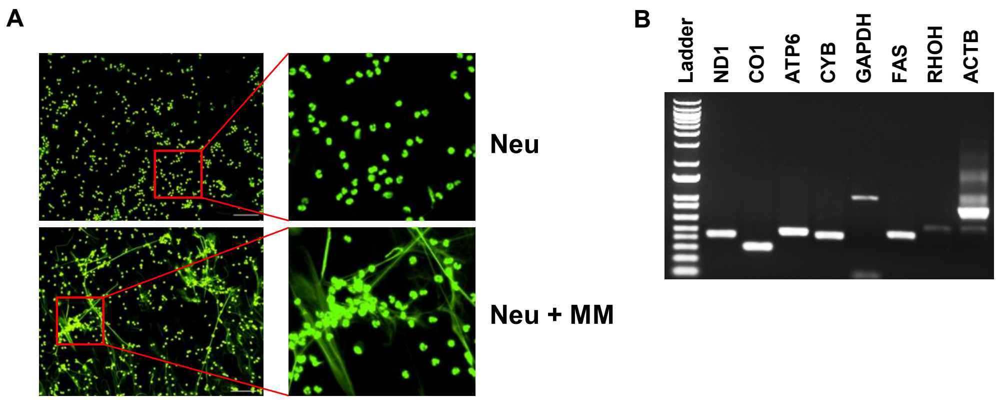


**Figure S3. MM cells induce NETosis, Related to Figure 3.** (A) Formation of NETs by neutrophils isolated from BM of MM patient cultured in the presence or absence of human MM cells evaluated by staining with Sytox Green nucleic acid stain. Representative images of NETs. Magnification 20x; scale bar, 50 µm. Insets, magnification 60x. (B) Presence of nuclear and/or mitochondrial DNA was determined by PCR using DNA isolated from supernatants collected from MM-stimulated human neutrophils. Ladder – 1 Kb Plus (ThermoFisher Scientific).

**Figure S4**

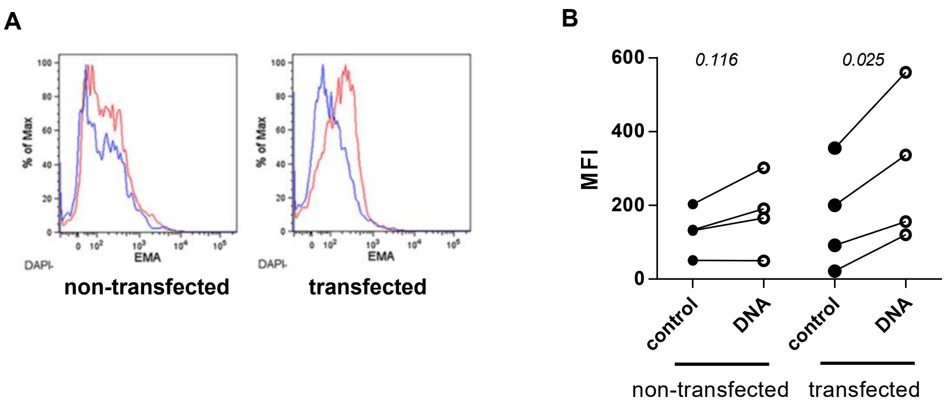


**Figure S4. Dendritic cells do not take up cfDNA, Related to Figure 5.** DCs were transfected with 1 µg of genomic neutrophil-derived EMA-labeled DNA. Uptake of DNA in transfected and non-transfected DCs was detected by flow cytometry 18h after the transfection. Representative flow cytometry plots (A) and individual values of MFI (B). Statistics: paired two-tailed Student's *t* test.

Figure S5

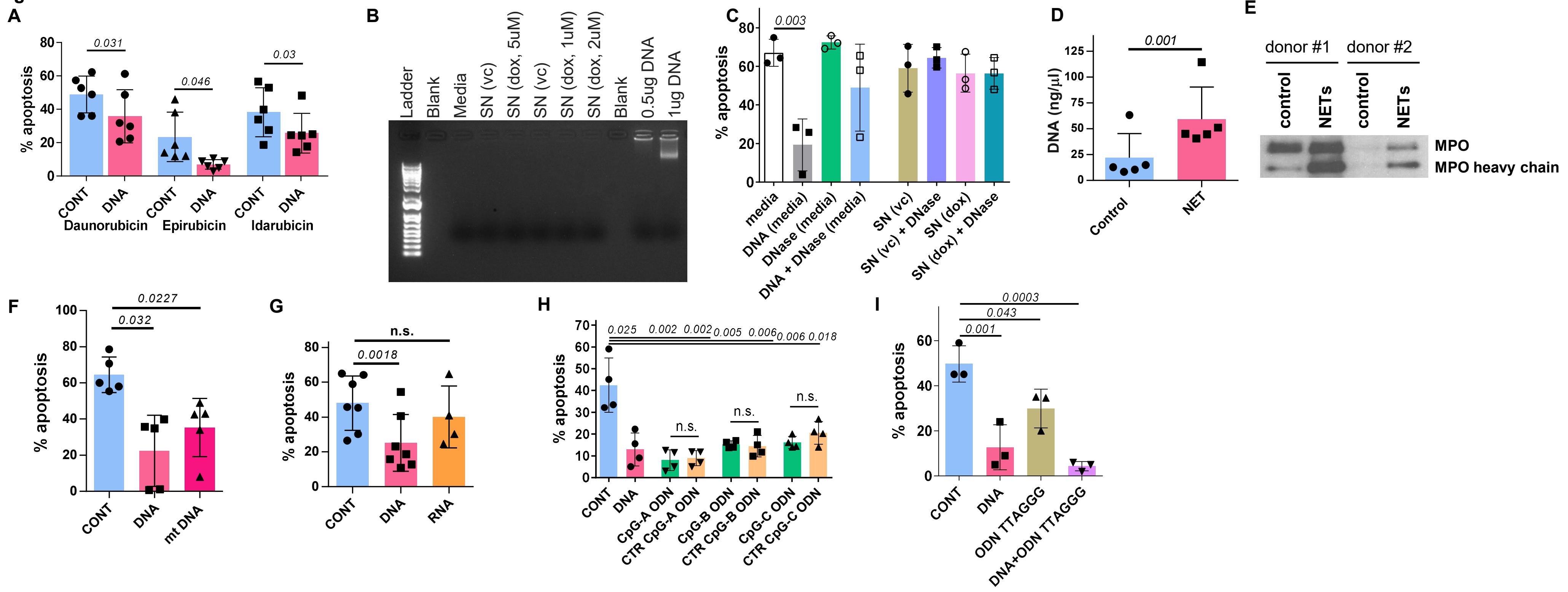


**Figure S5. Cell-free DNA protects tumor cells from anthracyclines, Related to Figure 5.** (A) Drug-specific apoptosis of U266 cells treated in the presence or absence of gDNA.
(B,C) DNA derived from apoptotic cells has no chemoprotective effect on tumor cells. H929 cells were treated for 1h with either doxorubicin (dox, 1-5µM) or vehicle control (vc, PBS) followed by washing away drug and culturing for an additional 24hr in drug-free medium. (B) Supernatants (SN) were then collected from vc-treated or dox-treated cells and run on agarose gel. Genomic DNA was used as a control. (C) Doxorubicin-induced apoptosis of H929 cells resuspended in these SN and treated with or without addition DNase I is shown. In control, MM cells were resuspended in culture medium (medium) and treated in the presence or absence of cfDNA with or without addition of DNase I.
(D,E) NETs or control neutrophil supernatant (control) were collected, respectively, from PMA-stimulated or unstimulated neutrophils from healthy donors. Cell-free DNA level (D) was measured and presence of MPO was detected by western blotting (E).
(F-G) Doxorubicin-specific apoptosis of MM1.S cells treated in the presence or absence of gDNA or mtDNA (F), RNA (G), CpG and control ODNs (H), TLR9 antagonist ODN or combination of cfDNA with TLR9 antagonist ODN (I).
Individual values, mean, and SD values for each experiment are shown. Statistics: paired 2-tailed Student's *t* test (A, C-G) and one-way ANOVA (H,I).

### Figure S6

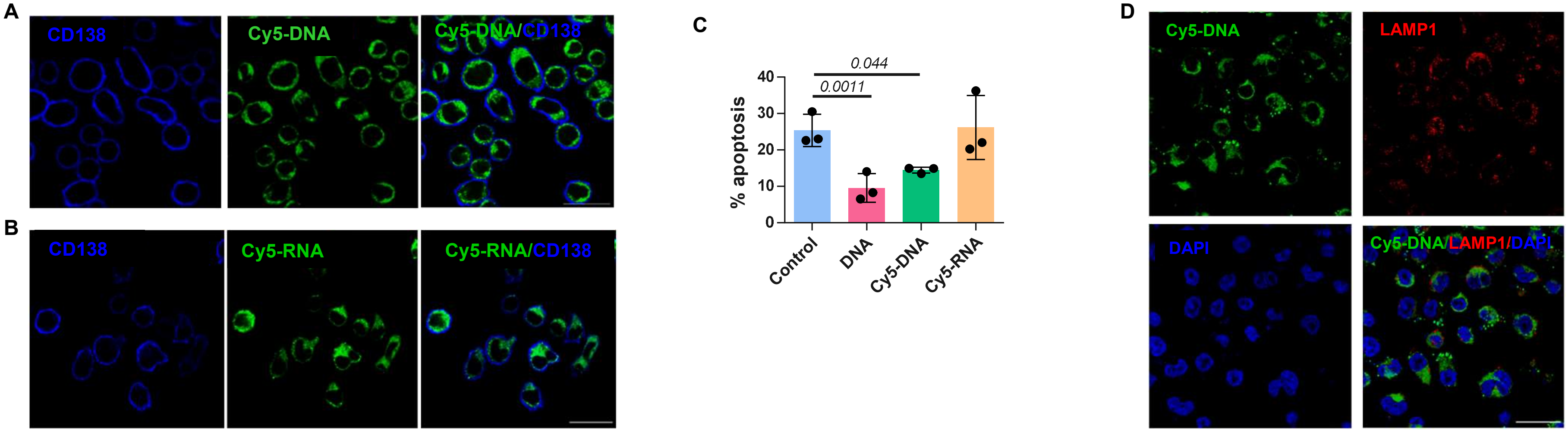


**Figure S6. Localization and effect of oligonucleotides on MM cells, Related to Figure 5.** (A,B) Representative images of Cy5-labeled DNA ODN (A) and Cy5-labeled RNA oligonucleotides (B) taken up by CD138+ MM1.S cells. Images were acquired using a confocal microscope. Scale bar, 20 µm. (C) Doxorubicin-induced apoptosis of H929 cells evaluated in the presence or absence of Cy5-DNA ODN and Cy5-RNA oligonucleotides. Individual results, mean and SD values are shown. Statistics: paired two-tailed Student's *t* test. (D) Intracellular localization of Cy5-labeled DNA ODN and LAMP1 in MM1.S cells was determined using confocal microscopy. Scale bar, 20 µm.

**Figure S7**

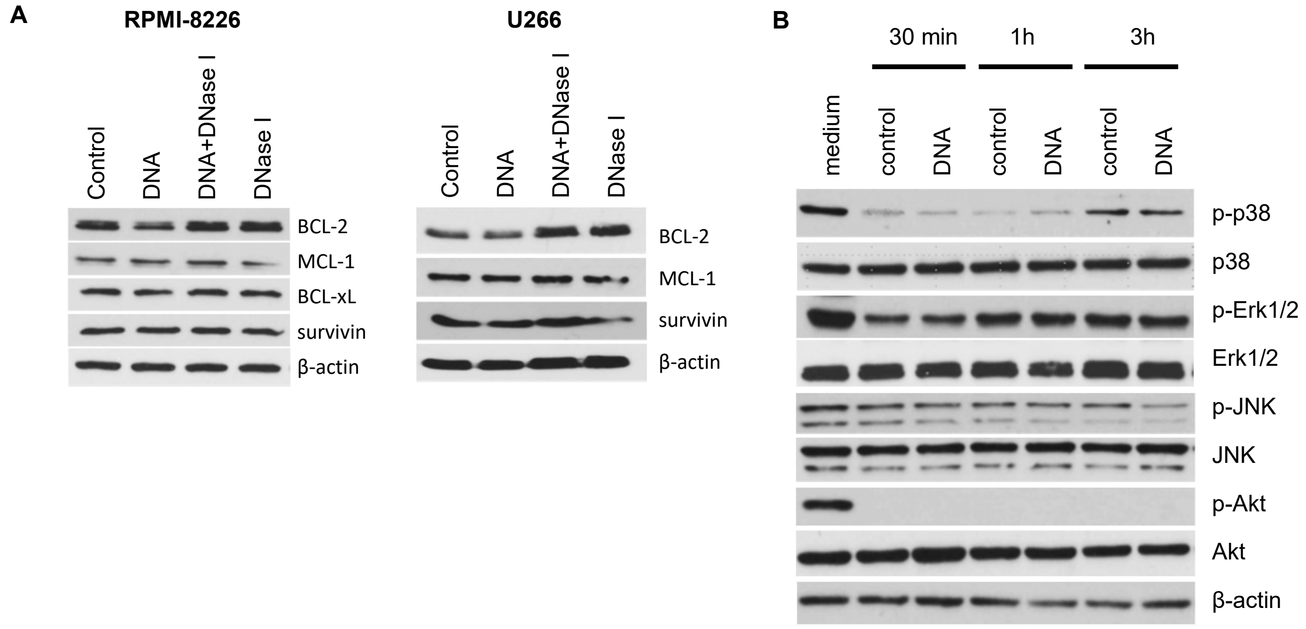


**Figure S7. Cell free DNA does not mediate pro-survival signaling, Related to Figure 5.** (A) Expression of anti-apoptotic bcl-2 family members was determined in MM cells incubated overnight with or without cfDNA. Representative western blots are shown. (B) RPMI-8226 cells were kept in serum-free medium for 2h followed by addition of DNA for indicated periods of time. Expression of phospho- and total proteins were detected by western blotting. Lane 1: cells were kept in medium with 10% FBS.
